# A 3-genes interferon signature predicts sustained complete remission in pediatric AML patients

**DOI:** 10.1101/2025.07.17.664572

**Authors:** Shimaa Sherif, Aesha Ali, Khadega A. Ibrahim, Darawan Rinchai, Mohammed Elanbari, Dhanya Kizhakayil, Mohammed Toufiq, Fazulur R. Vempalli, Tommaso Mina, Patrizia Comoli, Kulsoom Ghias, Zehra Fadoo, Sheanna Herrera, Che-Ann Lachica, Enas D.K. Dawoud, Hani Bibawi, Sandra Sapia, Blessing Dason, Anila Ejaz, Mohammed Y. S. Anas, Ayman Saleh, Giusy Gentilcore, Davide Bedognetti, Chiara Cugno, Sara Deola

## Abstract

The immunological composition of the microenvironment has shown relevance for diagnosis, prognosis, and therapy in solid tumors but remains underexplored in acute leukemias. We investigated the significance of acute myeloid leukemia (AML) bone marrow microenvironment in predicting chemosensitivity and long-term remission in pediatric patients.

We analyzed 32 non-promyelocytic pediatric AML patients at diagnosis using NanoString PanCancer IO 360 assay, RNA sequencing, and deep-phenotype flow cytometry analyses. The findings were validated using the pediatric TARGET AML dataset.

A short signature of three interferon (IFN)-related genes (*GBP1, PARP12, TRAT1*) distinguished patients with chemosensitive disease and reduced minimal residual disease after induction chemotherapy. The signature stratified patients overall, and within the clinically defined “standard-risk” group, patients with high gene expression at diagnosis had significantly longer overall survival. The leukemia microenvironment associated with this signature showed enrichment of non-exhausted CD4^+^ and CD8^+^ T cytotoxic lymphocytes and expansion of CD8^+^ T effector memory cells re-expressing CD45RA (TEMRA) in patients with a favorable prognosis. Our results show the importance of the bone marrow microenvironment in pediatric AML and provide tools for a refined stratification of “standard-risk” patients, lacking adequate risk-oriented therapies. They also offer a promising guide for tackling immune pathways and exploiting immune-targeted therapies.

**Statement of significance:** We identified a novel three-gene IFN-related signature that distinguished pediatric AML patients by chemosensitivity and remission outcomes. It stratified patients across all risk groups, including the “standard-risk” group, with high expression linked to a T-cell-enriched microenvironment and longer survival. This signature may enhance risk stratification and guide targeted immunotherapy.

## Introduction

Pediatric acute myeloid leukemia (AML) represents a complex and heterogeneous group of hematological malignancies that account for 20% of all pediatric leukemias. Despite remarkable breakthroughs in therapeutic approaches in the last decades, AML is still characterized by suboptimal outcomes, with an overall survival (OS) rate of approximately 65% (1, 2). Recent improvements in disease-free survival and risk of relapse have been achieved by the addition of a targeted tyrosine kinase inhibitor in patients with FLT3/ITD+ (3).

Numerous studies have emphasized the critical role of immune cell composition and behavior in the tumor microenvironment (TME) of solid cancers (4, 5). The contexture, functional orientation, and intricate interactions of immune cell subsets and tumor cells within the TME can directly influence patients’ treatment response and clinical outcomes (6-8).

Although it is tempting to speculate that the same model may be applicable to acute leukemia to exploit novel immunotherapies and extend patient survival (9-11), the role of TME in leukemia (L-TME) is still underexplored (12). The architecture of the bone marrow (BM) sustains normal hematopoiesis through delicate interactions in microenvironment subcompartments (13). Within BM niches, resident immune cells such as antigen-presenting cells, B cells, and memory T cells allow cognate antigen interactions and exchange immune information with the peripheral blood (PB) (14). At the onset of leukemia, the homeostasis of these fine-tuned networks is disrupted by the rapid invasion of leukemic cells, leaving a limited healthy counterpart available for assessment.

To increase the likelihood of meaningful immune network analysis in this scenario, we performed a gene expression study starting from an immune-directed platform, the NanoString PanCancer immune-oncology assay.

We explored the importance of the leukemic BM microenvironment (L-TME) in predicting chemosensitivity and ≥ 6 months remission in a multi-center group of 32 non-promyelocytic pediatric AML patients and validated the findings using data from the Therapeutically Applicable Research to Generate Effective Treatments (TARGET) AML database. Finally, we confirmed our results using 20-color flow cytometry analysis of the L-TME lymphoid composition on all available live-frozen BM samples in the cohort (nine samples).

## Methods

### Ethical compliance

The study was conducted in accordance with the ethical approval granted at all collaborative sites: Qatar, Sidra Medicine Institutional Review Board (IRB: #20110003636/2011; Pakistan, Aga Khan University Ethics Review Committee# 3825-Onc-ERC-15; Italy, IRCCS Fondazione Policlinico San Matteo Ethics Review Committee #1500786 and 1500787. All samples were collected after obtaining informed consent as appropriate.

### Clinical samples and cohorts

Thirty-two pediatric patients recruited in Qatar, Italy, and Pakistan provided BM samples for AML diagnosis. The patients were stratified and treated according to the COG AAML0531 protocol (15). Samples were live-frozen after Ficoll-gradient enrichment and channeled to Sidra Medicine for downstream analyses. Clinical and follow-up data for each patient were recorded, as detailed in Supplementary Table 1.

Of the 32 patients, 25 were analyzed using NanoString, and formed the Discovery Cohort. A subset of 19 patients from the Discovery Cohort, from whom sufficient RNA was available, was subsequently studied using mRNA-Seq for technical validation (Cross-Validation Subset). Next, an independent small cohort of seven newcomer patients was assessed using RNA-Seq, and an Internal Validation Cohort was formed. For pathway and gene enrichment analyses, all RNA-Seq cohorts (RNA Cross-Validation Subset, *n=19*; and Internal Validation Cohort, *n=7*) were combined in the “whole-RNA-Seq” cohort (Supplementary Table 1).

Patients achieving a 6-month complete remission (CR) after the first line of chemotherapy were assigned to “sustained-CR” group while patients who were refractory, died or relapsed within 6 months and/or obtained CR after allogeneic hematopoietic stem cell transplantation (allo-HSCT) were assigned to “non-sustained CR” group.

Five patients from the “non-sustained CR” group and 4 patients from the “sustained-CR” group were analyzed using a 20-color panel by flow cytometry (Deep Phenotype Cohort).

The Therapeutically Applicable Research to Generate Effective Treatments (TARGET) pediatric AML RNA-Seq database AAML1031 (3) was used as the External Validation Cohort. Patients with BM samples at diagnosis and complete clinical data available (*n*=833) were selected and assigned to “sustained-“and “non-sustained CR” groups with the same criteria applied to the cohort of 32 patients (Supplementary Table 2). To harmonize the risk stratification category nomenclature across different protocols, we use in the manuscript “standard” risk, referring to standard-or intermediate-risk patients.

### RNA analysis

BM samples were analyzed for RNA expression using the nCounter NanoString PanCancer immune-oncology IO 360 assay, which consists of a 770-gene panel enriched for immune-related genes. mRNA-sequencing (mRNA-seq) was performed at a depth of 20-million-reads on an Illumina HiSeq 4000 sequencer (Illumina, USA). Data analysis from fastq to the raw count generation was performed using the bcbio-nextgen (v1.2.0) rnaseq pipeline. The preliminary quality of sequencing reads was assessed using FASTQC (v.0.11.8). Raw reads were mapped to the human genome GRCh38.p13 (Genome Reference Consortium Human Build 38, INSDC Assembly GCA_000001405.28, Dec 2013) using STAR_2.6.1d aligner and featureCounts v2.0.0 was used to generate the raw counts.

Gene expression normalization for all cohorts was performed using the Variance Stabilizing Transformation (VST) of the DESeq2 R package within lanes to correct for gene-specific effects (including GC content) and between lanes to correct for sample-related differences (including sequencing depth) using EDASeq (v. 2.30.0).

### Flow-cytometry analysis

Patients with available live frozen BM cells (sustained CR, *n=4*; non-sustained CR, *n=5*, as described in Supplementary Table 1) were analyzed using a 20-color flow cytometry panel to characterize the infiltration of T, B, and NK cells in the L-TME. The cell number and viability were determined using a NucleoCounter® NC-200™ (ChemoMetec, Denmark). Cells were stained with Zombie UV Fixable Viability Dye (BioLegend, Cat#423108), treated with human Fc receptor block (BD Pharmingen, Cat# 564219) and mouse serum (Invitrogen, Cat#01-6501), stained according to the manufacturer’s instructions with antibody cocktails (Supplementary Table 3) extracellularly and intracellularly (FIX & PERM Cell Kit, Invitrogen Cat# GAS003), and acquired on a FACSymphony A5 (BD Biosciences, USA). Data analysis was conducted using the OMIQ software (Dotmatics: www.omiq.ai, www.dotmatics.com).

### Statistical analysis

All analyses, unless otherwise specified, were performed in “R” version 4.2.1. DEGs were analyzed using a log2 normalized expression matrix using limma (v. 3.52.4) (16), unadjusted *p*<0.05, and visualized with ggplot2 (v. 3.4.2) (17) and ComplexHeatmap (v. 2.12.1) (18).

Survival analysis was conducted using Survminer (v.0.4.9) (19) to generate Kaplan–Meier curves. Hazard ratios (HRs) and corresponding *p*-values were calculated by Cox proportional hazard regression analysis using the R survival package (v.3.5.7) (20). The overall *p*-value for comparing survival among the groups was determined using the log-rank test. Clinical variables included age at diagnosis (ordinal), clinical risk groups (categorical) and 3-genes signature enrichment divided into quartiles (categorical).

For pathway enrichment analysis, the DEGs in the CR groups (*p*<0.05) from the Whole-RNA-Seq combined cohort (19+7 samples) were uploaded to Ingenuity Pathways Analysis (IPA). Raw data were downloaded into R and plotted using the R package ggplot2 (v. 3.4.2).

Single-Sample-Gene-Set Enrichment Analysis (ssGSEA) was performed using normalized log2 expression data to calculate gene enrichment scores (ES) using the R package GSVA (v. 1.44.5) (21). Gene sets of immune cell-specific signatures were used as described by Bindea *et al*. (22) with slight modifications (23). ES were visualized using the ComplexHeatmap R package (v. 2.12.1) (18). Correlations were analyzed using Stat _cor in the ggpubr R package (v. 0.6.0). Statistical analyses of the flow cytometry data were performed using GraphPad Prism (v. 10.3.1).

## Results

### Patients with sustained-CR present a TH1-enriched L-TME at AML diagnosis

We performed DEG analysis using the NanoString assay on the Discovery Cohort between patients with sustained-CR (*n=6*) vs non-sustained CR (*n=19*), and identified a clear distinction between myeloid suppression-related genes and a T-cell infiltration signature represented by 67 DEGs (*p* ≤ 0.05) (Figure 1a) (Supplementary Table 4). By mapping these genes in the two CR groups, the T-cell signature, including cytotoxic T-cell infiltration (*CD3D, CD8A, PRF1, GNLY*), interferon (IFN) signaling activation (*IFIT3, IFITM1, IFI27, STAT1, GBP1, PARP12, TRAT1*), antigen presentation, and B-cell functions (*TAP1* and *CD19*), characterized patients with sustained-CR, while early loss of CR and refractory disease were marked by a pro-inflammatory response of macrophage-myeloid suppression signature (*CD68, TREM1*) and an intrinsic oncogene signaling linked to T-cell exclusion and immune suppression (*WNT5A, WNT*-*β*-catenin pathway, and *EFGR*) (Figure 1b).

**Figure 1.**
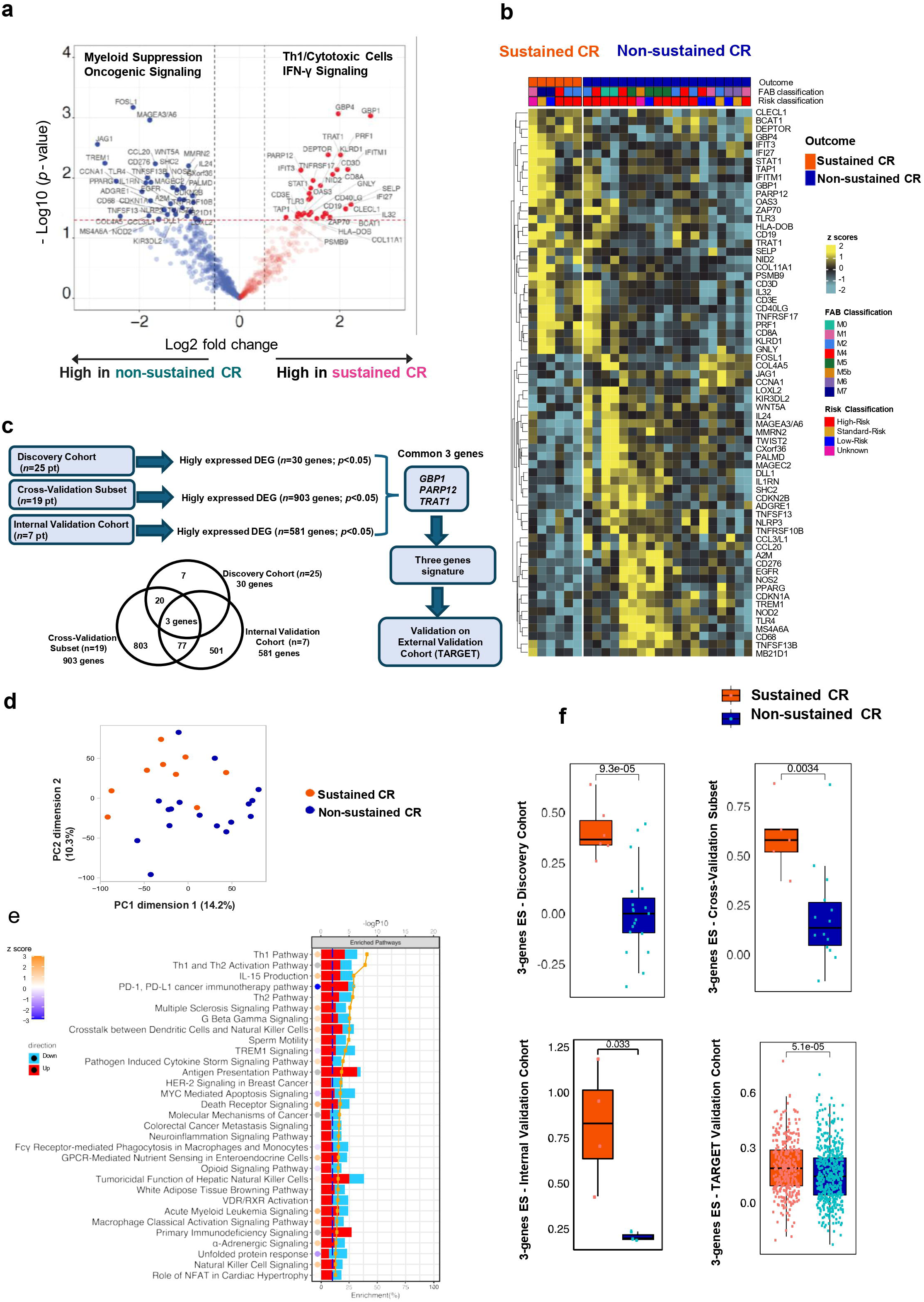
Identification of prognostic signature from the gene expression data of the Discovery Cohort, Cross-Validation Subset, and Internal Validation Cohort. (a) Volcano plot of differentially expressed genes in sustained-CR *vs* non-sustained-CR from the Discovery Cohort (*n*=25). Red dots:⍰high-expressed genes in the CR sustained groups (*p*-value⍰<⍰0.05, Fold Change of 1) and blue dots:⍰low-expressed genes in the CR sustained groups (p-value⍰< ⍰ 0.05, Fold Change of 1). (b) Heatmap showing the expression of 67 DEGs between the sustained-CR and non-sustained CR from the Discovery Cohort (*n*=25). FAB classification and clinical risk stratification are annotated on top of the heatmap. (c) A chart and Venn diagram showing the overlap among the highly expressed genes from the Discovery Cohort (*n*=25), the Cross-Validation subset (*n*= 19) and the Internal Validation Cohort (*n*=7). The intersection of these cohorts resulted in three common genes (*GBP1, PARP12, TRAT1*). The selection was performed by shortlisting significant DEGs (*p* <0.05) overexpressed in the sustained-CR group across all the cohorts. (d) PCA plot of the normalized expression values of the 26 patients from the whole mRNA-Seq cohort annotated by CR outcome. (e) IPA pathways analysis of DEGs (*p* <0.05) between the sustained-CR and non-sustained CR from Whole mRNA-Seq cohort (*n*=26). Red: upregulated in sustained-CR, blue, downregulated in sustained-CR. The orange line indicates the –log (*p*-value). The dots display the predicted activation state of the implicated biological functions reflected by the activation z score. The bases of this inferred activation state are literature-derived relationships between genes and the corresponding biological function. Pathways that are activated are marked with orange dots, indicating a positive activation score, whereas pathways inhibited are marked with a blue dot, indicating a negative activation score. A gray circle indicates that no literature-derived information exists to estimate the activation state. (f) Boxplots displaying the enrichment scores (ES) for the 3-genes signature between the sustained-CR and non-sustained CR in the different cohorts. *P*-value is calculated by Student’s *t*-test, and the middle black line represents the median.

To validate these results, we performed DEGs analysis using mRNA-Seq in a subset of 19 patients from the Discovery Cohort (Cross-Validation Subset), obtaining a total of 903 DEGs were overexpressed in the sustained-CR cohort, overlapping with the previous analysis. We then replicated the analysis in the Internal Validation Cohort, obtaining 581 overexpressed genes in the sustained-CR cohort (Figure 1c, DEGs lists are available in Supplementary Tables 5-6). Finally, we combined the mRNA-Seq-based cohorts (Whole mRNA-Seq Cohort, see Supplementary Table 1) to perform functional pathway analyses on the DEGs in patients with sustained-vs non-sustained-CR patients (Supplementary Table 7).

By applying Principal Component Analysis (PCA) to the transcriptome of the Whole-mRNA-Seq Cohort, we observed a separation pattern between sustained and non-sustained-CR (Figure 1d), suggesting biological differences between the compared groups. IPA enrichment pathway analysis confirmed that patients with sustained-CR benefit from an immune-rich L-TME, with a prevalence of T helper 1 (TH1) activation pathways, IL-15 production, and inhibition of PD1-PDL1 and myeloid TREM1 pathways, demonstrating an association between a TH1-skewed/cytotoxic L-TME and long-term remission (Figure 1e, Supplementary Table 8).

### A TH1-enriched L-TME at AML diagnosis is represented by a 3-gene IFN signature

To restrict this gene signature to a robust and clinically applicable one, we shortlisted the genes that were 1) common across all cohorts and 2) most highly expressed in DEG analysis. We identified three genes: guanylate-binding protein (*GBP1*), poly-ADP-ribose-polymerase-12 enzyme (*PARP12*), and T-cell receptor-associated transmembrane adapter (*TRAT1*) (Figure 1c). Notably, all three genes are related to IFN: *GBP1* (24, 25) and *PARP12* (26) are type II and type I IFN-stimulated genes innate immunity enhancers with strong antimicrobial properties, while TRAT1 is involved in cytotoxic/IFNγ-polarized immune responses (27) and correlates with immune cell infiltration and favorable prognosis in non-small cell lung carcinomas and diffuse large B cell lymphoma (27, 28).

Finally, we validated the identified IFN-signature in the TARGET database to evaluate its potential clinical relevance.

In the TARGET Cohort, the 3-genes-IFN signature successfully discriminated between patients with sustained and non-sustained-CR (Figure 1f), and performed better than all other genes in identifying patients with the lowest Hazard Ratio (HR) (Supplementary Figure 1a).

### The L-TME of patients with the 3-genes-IFN signature is enriched of non-exhausted CD4^+^ and CD8^+^ T cytotoxic lymphocytes

To explore the biological context behind our findings, we analyzed the L-TME of all available live-cryopreserved BM samples (*n=9*; 5 non-sustained-CR and 4 sustained-CR) in our cohort using a 20-colors lymphoid-skewed antibody panel (Supplementary Table 3). The CD45^+^ healthy L-TME portion was comparable between sustained- and non-sustained-CR samples (97.2±2% vs 85.4±9.1%, respectively). However, the cellular composition was strongly skewed, with a higher frequency of CD3^+^ T-lymphocytes in the sustained-CR (80.1±8.3%) compared to the non-sustained-CR group (14.1±12.2%) (Figure 2a and bI-cI), and a decrease in the myeloid healthy CD3^−^/CD19^−^ compartment (9.4±6% in sustained-CR and 63.7±32.5% in non-sustained-CR (Figure 2bI and bIII). A healthy age-matched BM to our cohort typically contains 50–70% myeloid cells, while lymphocyte levels range from 10-30% (29), suggesting that lymphocytes in the sustained-CR cohort actively expanded in the presence of leukemia blasts at the expense of the myeloid cell population. The % of blasts reported at the time of clinical AML diagnosis (combined microscope and flow cytometry analyses) was also lower in the sustained-CR cohort (Table 1, Figure 2bIV). A deeper analysis of the T lymphocyte subpopulations revealed that the frequencies of CD8^+^ and CD4^+^ cells were comparable between the two groups (Figure 2a and bII). Curiously, a small population of CD4^dim^/CD8^+^ or ^−^ cells characterized more non-sustained remission samples (Figure 2cI-II), representing IFNγ^+^ doublet T-cells engaged in close contact with other CD8^+^ and CD4^+^ cells, likely in cytolytic activity (see also Supplementary Figure 1b). The relative expression of IFNγ and CD107 was similar between the two cohorts, trending higher in non-sustained-CR CD4^+^ cells (Figure 3aI and bI); however, the cell composition between the cohorts showed remarkable differences. First, CD8 T effector memory cells (CD8^+^/CD62L^−^/CD45RA^−^ TEM) were significantly enriched in the non-sustained remission cohort, whereas CD8 T effector memory cells re-expressing CD45RA (CD8^+^/CD62L^−^/CD45RA^+^; TEMRA) were more prevalent in the sustained-remission cohort (Figure 3aII and bII-III). Second, TEM cells in the non-sustained remission cohort showed a larger FSC and a more exhausted phenotype, with PD-1 expression being significantly higher in the CD4^+^ compartment. Other exhaustion markers (TIM-3 and LAG-3) also trended towards higher levels than those in the sustained remission group (Figure 3a IV-VI and b IV-V).

**Figure 2.**
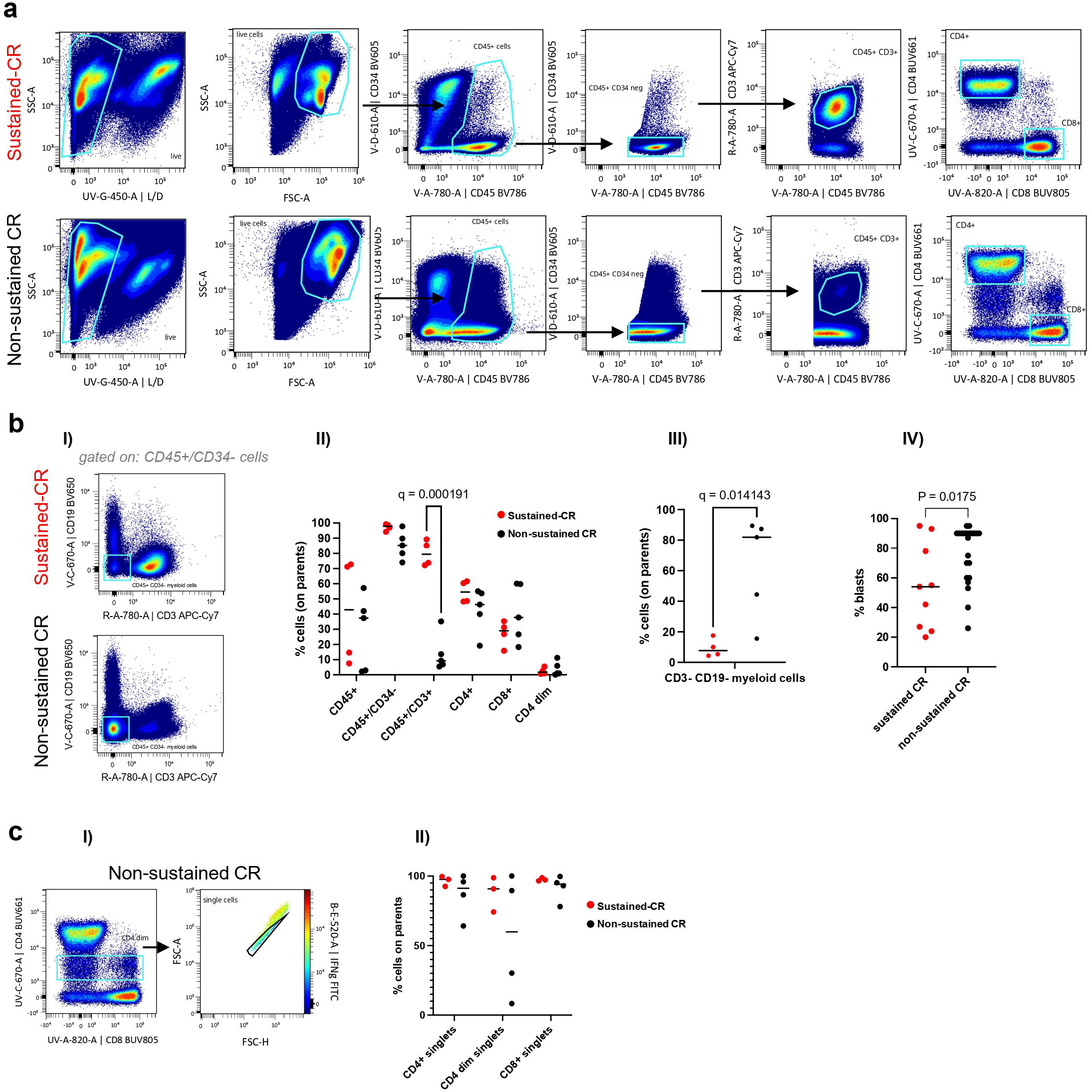
Lymphocyte profiling of the L-TME in pediatric AML patients. (a) Logical gating strategy showing the T cell composition of L-TME in AML patients with (top row, *n=4*) and without (bottom row, *n=5*) sustained-CR. The data are displayed cumulatively in the dot plots and individually in the graphs (c and e). The same cumulative and individual representation is depicted in all figures showing flow-cytometry data. (b) I) Cumulative plots displaying CD45^+^/CD34^−^/CD19^−^ and CD3^−^ myeloid cells in AML patients with (top row, *n=4*) and without (bottom row, *n=5*) sustained-CR. II-III) Individual values of samples in (a) and (b) are displayed in a graph; Q value is the result of FDR-corrected multiple t-tests between sustained (red) and non-sustained (black) patients’ samples. Only significant differences are shown. IV) The % of blasts scored in the patients’ diagnosis for all the cohort is displayed in the graph for patients with and without sustained-CR. The blast score was calculated by a combination of microscope and clinical flow-cytometry analyses and provided as clinical report for patients in the Pathology Laboratory of Sidra. (c) I) Phenotype of the CD4 ^dim^ cell population: the majority of the cells are IFNγ^+^ (shown in the colored Z-axis) CD4^+^ and CD8^+^ doublets, engaging in close networking interactions. A full panel of expression markers for this population is shown in Supplementary Figure 1b. II) Individual values of samples in (b) show a trend of enrichment in CD4^dim^ doublets (considering lower singlets) in patients without long-term remission.

**Figure 3.**
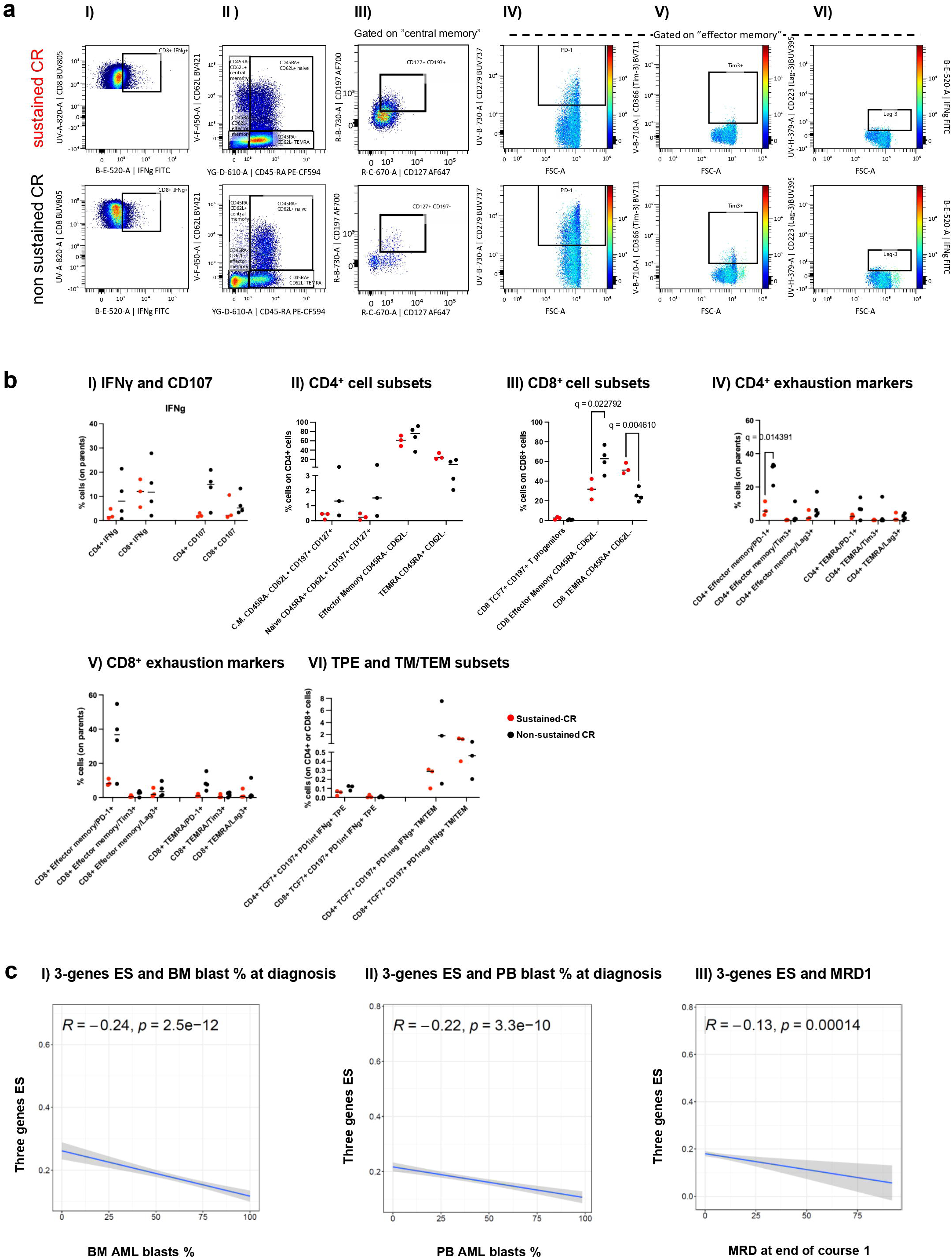
Characterization of lymphocyte subsets in the L-TME of pediatric AML patients. (a) Logical gating strategy, showing the measurement parameters of CD8^+^ cells (same logical gating was used for CD4^+^ cells) I) IFNγ positive cells, II) “central memory”, “naïve”, “effectors” and “TEMRA” cells, III) gating strategy to calculate naïve and central memory cells and IV-VI) PD-1, TIM-3, and LAG-3 exhaustion markers gated on “effector memory” cells in sustained and non-sustained-CR patients. The data are displayed cumulatively. Plots in IV-VI) show IFNγ in the colored Z-axis. (b) I-V) Individual values of samples, measured with the gating strategy in (a) are displayed in graphs. Some samples were excluded from the statistical analyses of subpopulations, due to insufficient number of acquired events, in these analyses we included (*n*=3) for sustained-CR, and (*n*=3-4) for non-sustained CR patients. Naïve and central memory cells were measured with the gating strategy showed in Figure 3a II-III selecting CD127^+^ (IL7Ra) and CD197^+^ (CCR7) on cells gated in II). VI) T Progenitor Exhausted (TPE) and T non-exhausted (TM/TEM) cells were measured according to criteria published in (30). Further gating strategies are shown in Supplementary Figures 2 and 3b. Q value is the result of FDR-corrected multiple *t*-tests between sustained (red) and non-sustained (black) patients’ samples. Only significant differences are shown. (c) Spearman correlations showing the relationship between the 3-gene ES score and the % of AML I) BM and II) PB blasts at AML diagnosis and III) with the burden of MRD measured after the first induction therapy (MRD1) in TARGET cohort.

We then assessed the L-TME T-cell memory compartment, according to a paired phenotype/TCR-functional classification recently described by Oliveira *et al.* in the context of melanoma (30). The paper described intra-tumor infiltrating memory T cells as: 1) primarily TCR-tumor specific but exhausted (T progenitor exhausted (TPE): TCF7^+^/CCR7^+^/PD-1^low/Int^/absent cytotoxic potential) and 2) viral/other non-primarily tumor TCR specific non-exhausted (T memory/T effector memory (TM/TEM): TCF7^+^/CCR7^+^/PD-1^−^/high effector cytokines). The frequencies of TPE and TM/TEM were similar in both groups, with a trend toward a higher presence in the non-sustained remission group (Figure 3bVI and Supplementary Figure 2). Finally, the presence of both CD4+ and CD8+ naïve and central memory cells, measured as CD62L^+^/CD45RA^+^/CD197^+^/CD127^+^ and CD62L^+^/CD45RA^−^/CD197^+^/CD127^+^, respectively, CD19^+^ B cells, (CD8^+^/CD3^−^/CD56^+^) NK, and (CD8^+^/CD3^+^/CD56^+^) NKT cells was minimal and similar in both groups (Figure 3aII-III, bII, and Supplementary Figure 3a-c).

Collectively, the phenotypic data provided evidence of functional differences in the L-TME of patients with different prognoses, showing a limited and more exhausted T-cell compartment in the L-TME of non-sustained-CR patients and a bulky-expanded less-exhausted T-cell compartment in patients with sustained-CR.

In this context, it is likely that our 3-genes IFN-related RNA signature captures a large abundance of functionally active T cells, approximately 6 folds higher (80% vs. 14% CD3^+^ infiltration) in patients with sustained remission.

### A high 3-gene-IFN signature enrichment at AML diagnosis is inversely proportional to AML leukemic burden

The lower number of blasts observed in the sustained-CR cohort suggests that a functionally T cell-enriched L-TME could contribute upfront (at AML onset) to contain the AML blast burden.

To test this hypothesis, we verified the relationship between the number of blasts in the BM and PB at AML diagnosis, and the 3-gene-IFN score in the TARGET cohort.

Both BM and PB blasts at diagnosis showed a clear inverse correlation with 3-genes ES (Figure 3c I-II). Although BM and PB blast numbers are not considered prognostic factors in pediatric non-promyelocytic AMLs (we confirmed no OS correlation with these variables in the TARGET cohort, data not shown), minimal residual disease after the first induction cycle (MRD1) is a clear biomarker of disease clearance and was found to have prognostic significance in the most recent WHO (adults) and COG AAML1831 (pediatric) classifications (31). We tested this variable in the ES and found an inverse correlation between the enrichment of the 3-genes-signature and the burden of MRD1 (Figure 3cIII).

These findings strengthen the evidence that a high 3-genes-ES represents a functional immune milieu at AML onset, contributing to disease eradication.

### A high 3-gene-IFN signature enrichment at AML diagnosis confers a longer OS to AML standard-risk patients

In COG AMML 0531 and 1031 protocols, the intensity of the treatment received by AML patients is calibrated on the relapse risk categorized as “low,” “standard” and “high” based on cytogenetic abnormalities and early response to treatment (15). The discovery and validation (TARGET) cohorts were then stratified accordingly.

Refined classifications integrating novel genomic data, including MRD1, are currently available for AML patients (31, 32).

However, standard-risk patients are in the “gray zone of a category formed by exclusion criteria from high- and low-risk definitions. An allo-HSCT intensified treatment is usually offered if an HLA-matched related donor is available (15, 33).

In the TARGET cohort, the OS of standard-risk patients’ OS is significantly lower than that of low-risk patients and comparable with that of the high-risk group that benefits from an intensified treatment (Figure 4a).

**Figure 4.**
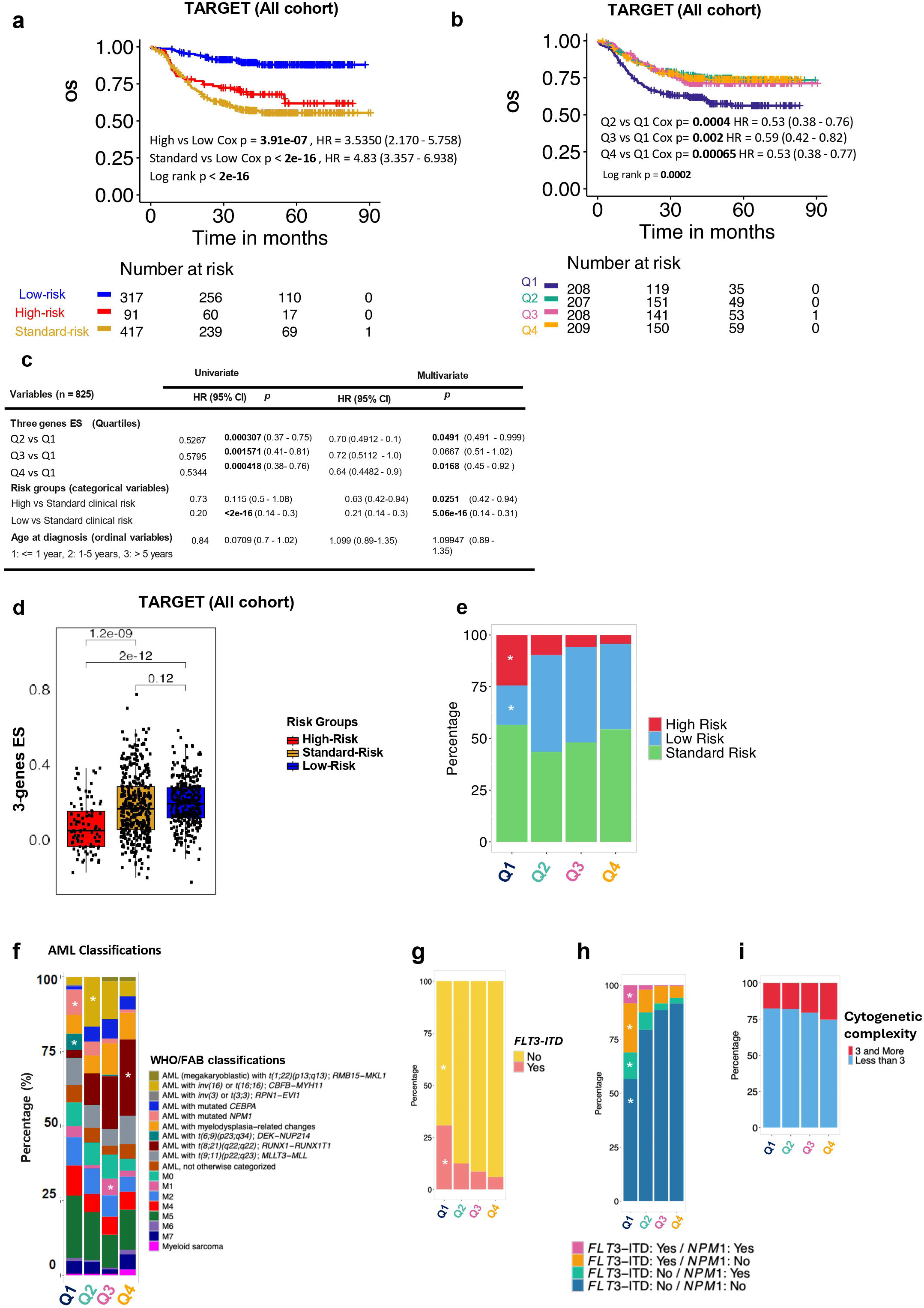
Prognostic value of the 3-gene IFN signature in pediatric AML OS. (a) Kaplan-Meier OS curves for clinical risk groups in the TARGET Cohort: low-, standard-, and high-risk groups. (b) Kaplan-Meier OS curve for the 3-genes enrichment score (ES) groups: The enrichment score was split into four quartiles within the entire TARGET cohort, with Q1 representing the lowest score and Q4 being the highest. (c) Univariate and multivariate analyses of OS in the TARGET cohort, evaluating the 3-gene ES quartiles, clinical risk-stratified groups, and age at diagnosis (*n*=825). (d) Boxplots showing the 3-genes ES across the clinical risk groups. *P*-values were calculated using the Student’s *t*-test, and the central black line represents the median. (e) Stacked bar chart showing the distribution of clinical risk groups across the quartiles of the ES signature in the TARGET cohort. White asterisks indicate*p*-value < 0.05, based on a chi-squared test for equal proportions (*n* =789). (f) Distribution of AML clinical subsets according to WHO and FAB classifications across the different 3-genes ES quartiles in the TARGET cohort. White asterisks indicate p-value < 0.05, based on a chi-squared test for equal proportions (n =784). A higher proportion of patients with AML *t(8;21)(q22;q22); RUNX1-RUNX1T1* was observed in the highest 3-genes ES quartile (Q4) than in the other ES groups (*p* = 7.83e^−10^). A higher proportion of patients with AML with mutated *NPM1* and *t(6;9)(p23;q34); DEK*-*NUP214* was present in the lowest 3-genes ES quartile (Q1) compared to other ES groups (p = 0.00031 and 4.282e^−06^ respectively). (g-h) Stacked bar charts showing: g) the distribution of *Flt3-ITD* mutations across the ES quartiles and h) the distribution of combined *Flt3-ITD* and *NPM1* mutations. The white asterisk indicates a *p*-value of 4.685e^−13^ based on a chi-square test for equal proportions of *FLT3-ITD* mutation status across ES quartiles (*n*=796), and a *p*-value < 2.2e^−16^ for combined *Flt3-ITD* and *NPM1* mutations status across score quartiles (*n*= 789). (i) Stacked bar chart showing the distribution of cytogenetic complexity groups within the quartiles of the ES signature in the TARGET Cohort (*n*=789). Patients with <3 cytogenetic alterations are shown in blue and those with ≥ 3 cytogenetic alterations are shown in red.

We performed a long-term (90-months) OS analysis of the TARGET Cohort according to the 3-IFN gene signature, quantified in “quartiles” enrichments and we discovered that the lowest ES “Q1” in the L-TME of patients was associated with a significantly lower OS (Figure 4b). To benchmark the importance of ES against clinically relevant parameters (cytogenetics, molecular mutations, and morphological remission after first induction), we performed univariate and multivariate analyses measuring the ES quartiles against the clinical-risk group stratification, which included all the relevant variables. The importance of the ES in determining the OS was maintained particularly in the extreme enrichments Q1 (lowest) vs Q4 (highest) in the multivariate analysis (Q4 vs Q1, p = 0.0168, Figure 4c).

Next, we analyzed the distribution of the 3-IFN gene signature across clinical risk groups. The signature was enriched in the low- and standard-clinical risk groups (Figure 4d). Accordingly, high-risk patients were enriched and low-risk patients were depleted of the less favorable Q1 ES signature, whereas the quartiles were evenly distributed across standard-risk patients (Figure 4e). When we analyzed the scores across the FAB and WHO AML classifications, we found that patients with low-risk *t(8;21)(q22;q22.1)/RUNX1:RUNX1T1* translocations were predominantly represented in the Q4 (high ES) group, whereas those with *NPM1* mutations and *t(6:9)(p23;q34)/DEK:NUP214* translocations were more prevalent in the unfavorable Q1 (low ES) group (Figure 4f). Next, we screened the distribution of scores of patients with *Flt3-ITD* mutations. Unfavorable Q1 quartile was more prevalent among *Flt3-ITD* mutated patients (Figure 4g). When *Flt3-ITD* was combined with *NPM1* mutation, Q1 remained predominant across all *Flt3-ITD/NPM1* combinations, whereas Q2-4 were enriched in patients without mutations (Figure 4h). Finally, we checked the quartiles distribution in patients with or without complex cytogenetics (≥3) and found that the quartiles were evenly distributed in these patients (Figure 4i).

Surprisingly, the ES was higher in infants (Figure 5a-b). A negative correlation between age and 3-genes ES was further confirmed by correlation analysis (Supplementary Figure 3d). This finding could not be assessed in our cohort, as only one patient was younger than 1-year old. Still, it aligns with recent literature suggesting that infant T cells may have innate-like functions and are prompt to respond to danger signals response (34-36). Indeed, the possibility of stratifying infant AML using a favorable prognostic factor could be promising, considering that this patient group usually has poor clinical outcomes (37).

**Figure 5.**
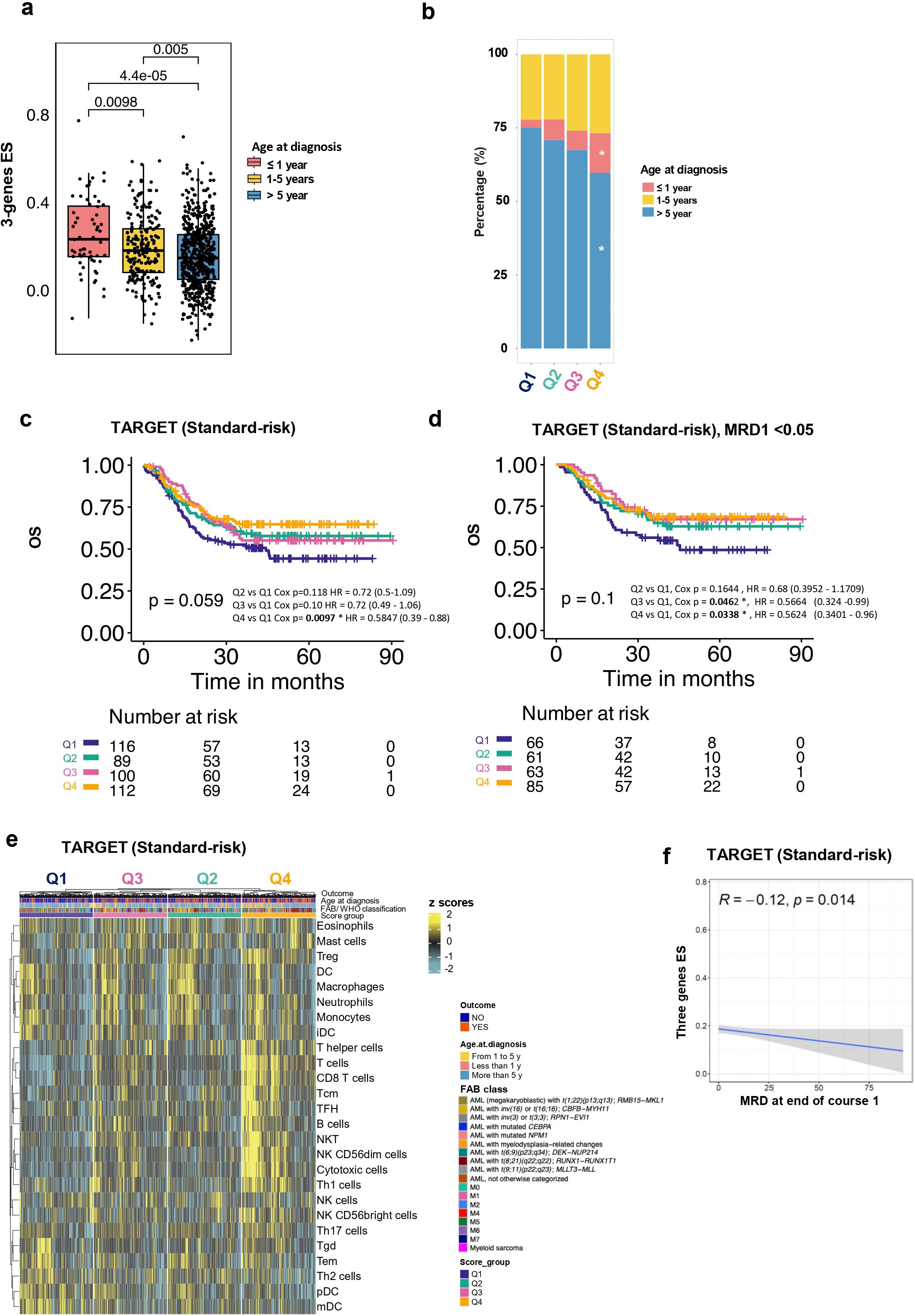
3-gene IFN signature and OS in standard-risk pediatric AML patients. (a) Boxplots showing the 3-genes ES across different age groups at diagnosis. *P*-value was calculated by Student’s *t*-test, and the central black line within each box represents the median. (b) Stacked bar chart showing the distribution of the variable “age at diagnosis” within the quartiles of the ES signature in TARGET Cohort. The chi-square pairwise proportion test showed the lowest proportion of the age group >5 years in patients with the highest 3-genes ES compared to other ES groups (p = 0.0058); and a higher proportion of age group ≤ 1 year in the same group (*p* = 0.00054). (c) Kaplan-Meier OS curve for the 3-genes ES groups split in quartiles in the clinical standard-risk group of the whole TARGET cohort. *P*-values of the compared quartiles, obtained with a Cox regression analysis are annotated in the graph (*n*=417). (d) Kaplan-Meier OS curve for the 3-genes ES groups split in quartiles in the clinical standard-risk group of TARGET cohort after removing all patients with MRD1 =/>0.05. P-values from Cox regression analyses comparing quartiles are annotated in the graph (*n*=275). (e) Heatmap displaying immune cells subpopulation enrichment scores across 3-genes ES groups within the standard clinical risk patients’ group. These immune signatures were previously described in Sherif *et al* (23). For each ES group CR outcome AML classification, outcome (sustained-/non-sustained CR) and age at diagnosis are annotated on top of the heatmap. (f) Spearman correlations showing the relationship between the 3-gene ES score and the burden of MRD measured after MRD1 in the clinical standard-risk group.

To investigate whether the 3-genes score could improve stratification within the risk groups, we tested the OS in each group. A survival advantage was granted by high 3-genes-ES only in the standard-risk cohort (Q1 vs Q4, *p* = 0.0097) (Figure 5c).

Specifically, a logistic regression analysis revealed that children with standard-risk AML and in the Q4 high 3-gene ES quartile had 100% increased odds of achieving long-term OS compared to those in the Q1 low ES quartile (Supplementary Figure 3e).

While we lacked sufficient genetic information to re-stratify patients according to the latest COG AAML1831 protocol and test the 3-gene ES in these new risk categories (32), we used MRD1 data to assign MRD1≥ 0.05% to high-risk following the newest recommendations. While the OS in all TARGET cohorts did not substantially change (Supplementary Figure 3f), the 3-genes ES in the standard-risk cohort still highlighted a significant negative impact on OS for patients with a low score (Q1) (Figure 5d).

To confirm that the 3-gene ES reflected the previously described cytotoxic-enriched L-TME in the standard-risk subjects of the TARGET cohort, we performed an enrichment analysis using a published immune subpopulations genes set (38). The analysis confirmed that the 3-genes-score represents a contextual cytotoxic/NK/Th1 expression profile at AML onset, depleted in Q1, and enriched in Q4 ES quartiles (Figure 5e).

Finally, we confirmed an inverse correlation between MRD1 and 3-gene-ES in this cohort (*p*= 0.014, Figure 5f).

## Discussion

The TME has been extensively described in the solid tumor literature: “hot” T-cell infiltrated tumors have been associated with a favorable prognosis, in contrast to “cold” tumors (23, 39).

While the leukemic composition of the AML TME has been extensively investigated, (40-42) the immune L-TME has been less studied. Recent literature describes how the TME is deeply affected by AML (13), but the correlation between a clear pattern of immune infiltration and prognosis remains controversial (43). Mazziotta et al. recently correlated a skewed trajectory of CD8 cells towards senescence with a worse prognosis in adult AML patients, analyzed with a multi-dimensional approach onset and relapse paired samples (44).

Our results, while aligning with their findings regarding the onset of pediatric AML leukemia, add important data on functional T cell infiltration and expand the analysis to include CD4^+^, CD19^+^, NK, and NKT cells.

Lasry et al (45) used single-cell RNA sequencing to profile sorted BM immune cells from 22 pediatric and 20 adult AML patients. They described an 11-genes inflammatory signature (iScore) that included IFN-related genes along with genes related to antigen presentation (HLA genes), growth factors signaling (HGF), and stress signaling oncogenic processes (HSP90AA1), which correlated with a worse prognosis. A key difference from our analysis, in addition to the single-cell approach, is the selection criteria of the pediatric TARGET validation cohort, which included 336 patients. In our study, we evaluated the iScore using whole RNA from the TARGET samples we tested (*n*=833); however, we could not confirm its association with worse prognosis (Supplementary Figure 3g I-III).

Our results seem to suggest a narrower IFN-rich functional signature from this broader inflammatory signature, representing a favorable L-TME. Deep flow cytometry analysis revealed that patients with this type of L-TME benefit from a 6-times higher abundance of CD3^+^ T cells enriched with TEMRA, a non-exhausted PD-1 negative phenotype, along with a lower number of infiltrating blasts. Consistently, patients exhibiting an enriched cytotoxic L-TME in the broad TARGET cohort displayed a lower number of blasts at diagnosis, achieved a lower MRD1, and increased long-term OS.

The ES was enriched in TARGET cohort patients assigned to low-and standard-clinical risks, and in general, a favorable high-3-genes score (Q4) aligned with good prognostic features, such as AML with *t(8;21)(q22;q22), RUNX1-RUNX1T1* and absence of Flt3-ITD mutations. Accordingly, the unfavorable 3-genes-depleted score (Q1) was associated with *Flt3-ITD* mutations and *t(6;9)(p23;q34); DEK-NUP214*, as well as with *NPM1* mutation in the absence of *Flt3-ITD*, per se a good prognostic factor.

It is tempting to assume that certain pediatric AML blast features are more immunogenic than others, triggering a favorable immune expansion at AML onset, making it more prone to control blast invasion, lowering the burden of MRD, and providing better survival.

Although our results point more towards an early clearance of blasts and we did not find any difference in CD4 and CD8 T stem cell memory enrichment between cohorts, it might also be possible that a pool of leukemic specific T-memory stem cells could better survive chemotherapy within a favorable T cell-rich L-TME, contributing to long-term AML disease control.

Such hypotheses are beyond the scope of this study and may be tested using a dedicated functional study in a larger cohort.

When measured within the clinical risk groups, the 3-genes score stratified the OS of standard-risk patients into four quartiles, showing a clear OS advantage for the highest ES quartile (Q4) and a 2-fold increased risk of having a reduced OS for patients with the lowest enrichment (Q1). Importantly, the ES was correlated with OS in standard-risk patients, even after removing patients with MRD1=/>0.05% (as per latest stratification guidelines) proving to be an independent risk-score that could offer to “Q1” patients the chance to improve their OS if assigned to an intensified treatment.

A limitation of this study is the small size of our Discovery Cohort, which is representative of three independent recruitment sites (Italy, Qatar, and Pakistan). Nevertheless, the analyses initially guided by the NanoString panel and focused on subsets of tumor-infiltrating genes likely favored a less-sparse L-TME analysis in the discovery phase.

Validation experiments using deep phenotype data were also based on a limited number of samples and did not allow meaningful correlations between leukemia genotype, phenotype, and T cell infiltration.

In summary, we identified a signature of 3-IFN-related genes, representing a non-exhausted T cell-enriched L-TME at leukemia onset, which correlates with better OS in pediatric AML patients.

Careful flow cytometry evaluation revealed the following characteristics of this favorable L-TME: 1) significant CD3^+^ T cell expansion with respect to the myeloid and malignant blast compartments (both cell types were not analyzed in detail in our work), 2) a relative increase in TEMRA CD8^+^ T cells, and 3) a less-exhausted phenotype.

This discovery, once tested in appropriate clinical trials, has the potential to enhance the stratification of standard-risk patients who currently lack appropriate risk-oriented treatment options. Furthermore, these findings may serve as a valuable roadmap for addressing immune pathways and exploring the potential efficacy of immune-targeted therapies.

## Declarations

## Acknowledgements

The authors thank the patients and their families for their participation in the study. We also acknowledge Sara Tomei and Lisa Sara Mathew for performing transcriptome analyses.

## Authorship

### Authors’ contributions

S. Sh., A. A., D. R., F. V., K. A. I., E. D. K. D., H. B., and S. Sa. and M. T. data curation and formal analysis; M. E. formal analysis and data visualization; T. M., P.C., K. G.,Z. F., B. D., A. E., M. Y. S. A., D. K., S. H., C. L., and A. S. provided resources and performed the investigation; S. D., S. Sh, A. A D. B. and C. C. conceptualization, validation, and writing. All the authors have reviewed the manuscript.

### Consent for publication

All authors have contributed to the manuscript and approved the submitted version. All the authors have reviewed the manuscript.

### Correspondence

Chiara Cugno, Research Department and Pediatric Hematology and Oncology Department, Sidra Medicine, Qatar, Doha.

## Ethics approval and consent to participate

Informed consent to participate in the study was obtained from all participants and parents or legal guardians of children under 16 years of age.

This Registry study was conducted with ethical approval from the Sidra Medicine Institutional Review Board (IRB) (protocol # IRB #20110003636/2011). This study was approved by the AKU Ethics Review Committee (approval number: 3825-Onc-ERC-15).

The approval numbers of the comitato Ethics Review Committee were # 1500786 and # 1500787.

## Availability of data and materials

The patients’ clinical data can be found in a data supplement available in the online version of this article. RNA expression matrices and TARGET clinical data were deposited at https://doi.org/10.6084/m9.figshare.24152742.

The R code used for this analysis can be found in the GitHub repository (https://github.com/Sidra-TBI-FCO/IFN-CR-AML) or in the Zenodo snapshot of this repository, created at the time of manuscript acceptance.

All the flow-cytometry data and analyses are available upon request.

---

**Supplementary Table 1**

Clinical and molecular characteristics of pediatric AML patients, including cohort and subcohort membership. The recruitment of pediatric patients involved data collection from 32 individuals diagnosed with AML at three distinct locations: Sidra Medicine in Doha, Qatar, IRCCS Fondazione Policlinico San Matteo in Pavia, Italy, and Aga Khan University in Karachi, Pakistan. Sustained-complete remission (CR) refers to patients who achieved a 6-month CR after first-line chemotherapy, while non-sustained CR included patients who had refractory disease, died or relapsed within 6 months, and/or achieved CR after allo-HSCT.

allo-HSCT, allogeneic hematopoietic stem cell transplantation; AML, acute myeloid leukemia; CR, complete remission; FAB, French-American-British classification; KAR, Karachi-Pakistan; PV, Pavia-Italy; SDR, Sidra Medicine-Qatar; WHO, World Health Organization.

**Supplementary Table 2**

The clinical and molecular characteristics of the TARGET pediatric AML RNA-Seq dataset (AAsML1031) were used as External Validation Cohort. Patients with available diagnostic BM samples and complete clinical information were included (*n*=833).

**Supplementary Table 3**

List of antibodies used for extracellular and intracellular staining for flow-cytometry analysis.

**Supplementary Table 4:**

List of 67 differentially expressed genes (DEGs) identified by NanoString analysis in the Discovery Cohort. Gene expression profiles were compared between patients with sustained-CR (*n*⍰=⍰6) and those with non-sustained CR (*n*⍰=⍰19). All DEGs met the significance threshold (*p*⍰<⍰0.05).

**Supplementary Table 5:**

DEGs in the sustained-CR group were identified by mRNA-Seq in the Cross-Validation Subset (n = 19). Gene expression profiles were compared between patients with sustained-CR (*n*⍰ =⍰5) and those with non-sustained CR (*n*⍰=⍰14). All DEGs met the significance threshold (*p*⍰<⍰0.05).

**Supplementary Table 6:**

DEGs in the sustained-CR group were identified using mRNA-Seq in the Internal Validation Cohort (*n*=7). Gene expression profiles were compared between patients with sustained-CR (*n*⍰=3) and those with non-sustained CR (*n*⍰=4). All DEGs met the significance threshold (*p*⍰<⍰0.05).

**Supplementary Table 7:**

DEGs in the sustained-CR group were identified by mRNA-Seq of the Whole mRNA-Seq Cohort (n=26). Gene expression profiles were compared between patients with sustained-CR (*n*⍰=8) and those with non-sustained CR (*n*⍰=18). All DEGs met the significance threshold (*p*⍰<⍰0.05).

**Supplementary Table 8:**

Functional pathways associated with DEGs between patients with sustained-CR (*n*⍰=8) vs non-sustained CR (*n*⍰=18) based on combined mRNA-Seq cohorts (Whole mRNA-Seq Cohort).

**Supplementary Figure 1:**

Prognostic HR of IFN signature and T-cell doublets in pediatric AML.

(a) Boxplot showing the Cox proportional hazard ratio (HR) for the 3 genes signature and genes with *p*-value < 0.05 (2766 genes) out of the 18000 genes of all RNA-Seq matrices in the standard-risk group of the TARGET Cohort. The 3-genes signature had the lowest HR among all the significant genes.

(b) Dot plots representing the cumulative rate of single cells, displayed as FSC-H vs FSC-A, across three T-cell populations: CD8^+^ CD4^+^ and CD8/CD4^dim^. In each set of three plots, the Z-axis represents a different antigen measured in patients with sustained-CR (red legend) and non-sustained CR (black legend). The analysis highlighted the increased doublet population within the CD4 ^dim^ gated cells, particularly positive for CD4, CD8, and IFN markers (indicated by the higher orange/red color intensity).

**Supplementary Figure 2:**

T-Cell subsets gating in BM of pediatric AML patients.

Dot plots representing the gating strategy used to analyze T Progenitor exhausted (TPE)and non-exhausted T memory/effector cells (TM/TEM), measured in CD8^+^ and CD4^+^ T-cells populations from patients with sustained (red) and non-sustained CR (black). The individual values are shown in Figure 3. On the left, the PD1 high/int/low gates are shown for the total CD4^+^ and CD8^+^ cell populations. In the middle, the selection of the TCF7 high cells, followed by logical gating measuring PD1 and IFNγ expression in these cells. TPE was defined as IFNγ neg/ PD1 int/low, whereas TM/TEM were identified as PD1 neg/IFNγ high. Cumulative data are shown here, indicating a higher presence of both TPE and TM/TEM cells in the non-sustained CR cohort, although individual differences were not statistically significant (Figure 3). Below are the control gates set up on the FMO samples using the same markers.

**Supplementary Figure 3:**

BM immune subsets and prognostic 3-Gene signature in pediatric AML

(a) Dot plots show the cumulative rates of B cells, NK cells, and NKT cells in patients with sustained (red) and non-sustained CR (black).

(b) Logical gating strategy used to identify CD4^+^ Naïve and Central Memory T cells.

(c) Individual values correspond to plots in (a) and (b) for patients with sustained-CR (red dots) and non-sustained CR (black dots).

(d) Pearson correlation between the 3-genes ES score and age at diagnosis in the TARGET cohort measured across three age groups: ≤ 1 year, 1-5 years and > 5 years.

(e) Logistic regression analysis showing the odds ratios with 95% confidence intervals (CI) of the 3-genes ES quartiles as an explanatory variable for CR in pediatric AML patients with clinical standard-risk in the TARGET cohort.

(f) Kaplan-Meier OS curve for clinical-risk groups in the TARGET cohort: low-, standard-, and high-risk groups after re-assignment of all patients with MRD1 ≥ 0.05% in high risk (*n*=789).

(g) Analysis of Lasry’s pediatric inflammation-associated gene score (iSCORE) in the TARGET AML cohort (*n* = 833): (I) Boxplots displaying the iSCORE defined by Lasry et al. (45), based on expression of 12 genes (*COTL1, GSN, HGF, HLA-DQA1, HLA-DQA2, IFI6, IFITM2, RHOG, HLA-DPA1, COMMD3, HSP90AA1*), compared between the sustained-CR and non-sustained CR patients. P-value was calculated using Student’s *t*-test; the central black line represents the median. (II) Boxplots showing iSCORE distributions across clinical risk groups in the TARGET cohort. *P*-values were calculated using the Student’s *t*-test, and the central black line represents the median. (III) Kaplan–Meier OS curve for the TARGET cohort stratified by iSCORE quartiles. The enrichment score was split into four quartiles, with Q1 representing the lowest iSCORE and Q4 being the highest score.

## Abbreviations

AML: Acute myeloid leukemia
allo-HSCT: Allogeneic hematopoietic stem cell transplantation
BM: Bone marrow
CR: Complete remission
DEGs: Differentially expressed genes
ES: Enrichment score
FAB: The French-American-British classification
GBP1: Guanylate-binding protein
IFN: Interferon
IPA: Ingenuity Pathway Analysis
ISGs: Interferon stimulated genes
L-TME: Leukemic bone marrow tumor microenvironment
MRD: minimal residual disease
OS: Overall survival
PARP12: Poly-ADP-ribose-polymerase-12 enzyme
PB: Peripheral blood
RNA-Seq/mRNA-Seq: mRNA-Sequencing
ssGSEA: Single Sample Gene Set Enrichment Analysis
TARGET: Therapeutically Applicable Research to Generate Effective Treatments
TH1: Thelper 1
TME: Tumor microenvironment
TRAT1: T-cell receptor-associated transmembrane adapter
tSNE: *t*-Distributed Stochastic Neighbor Embedding
WHO: World health organization

