## Supplementary figures and images for "A 3-genes interferon signature predicts sustained complete remission in pediatric AML patients"

### Supplementary Figure 1.jpg

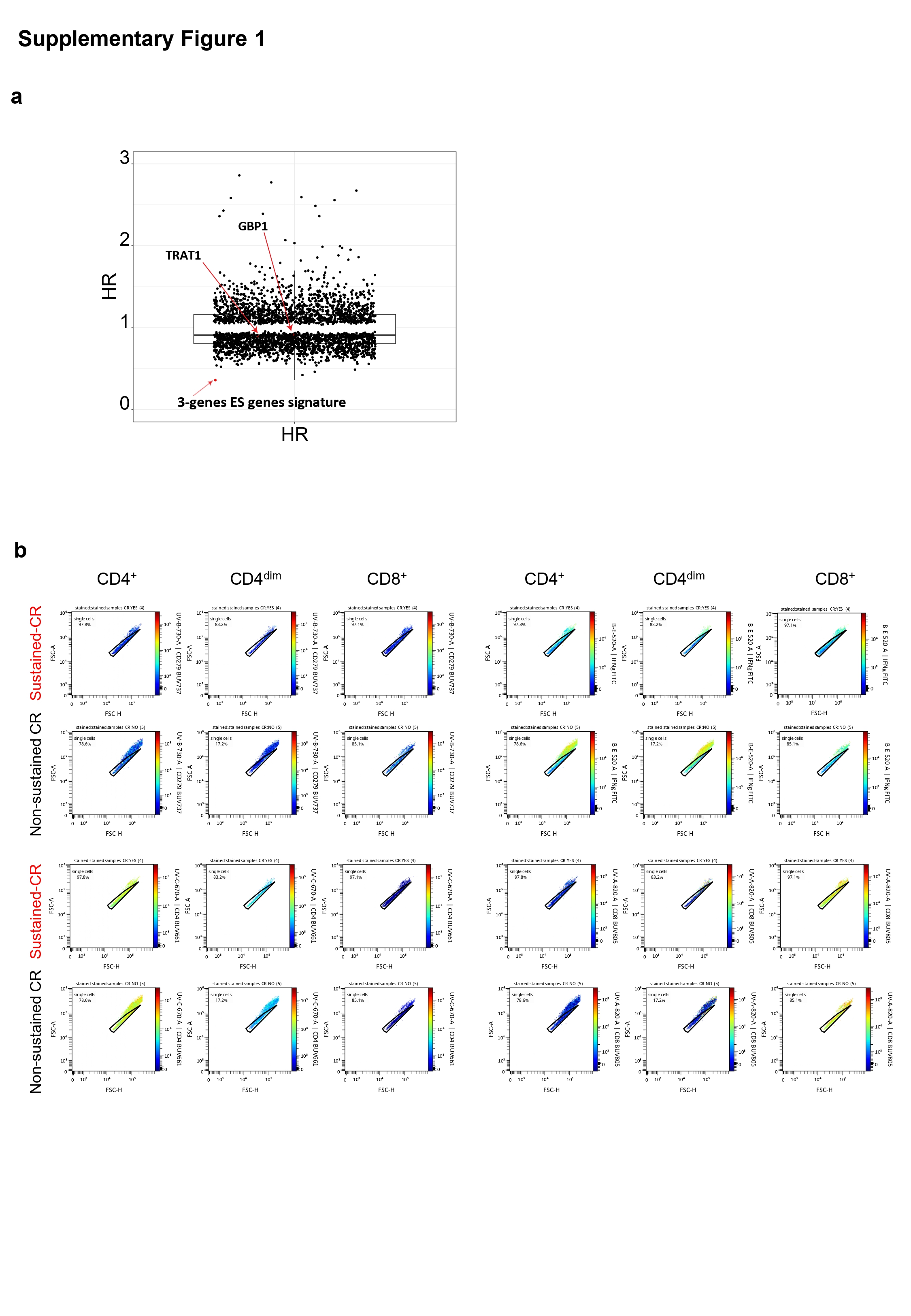

### Supplementary Figure 2.jpg

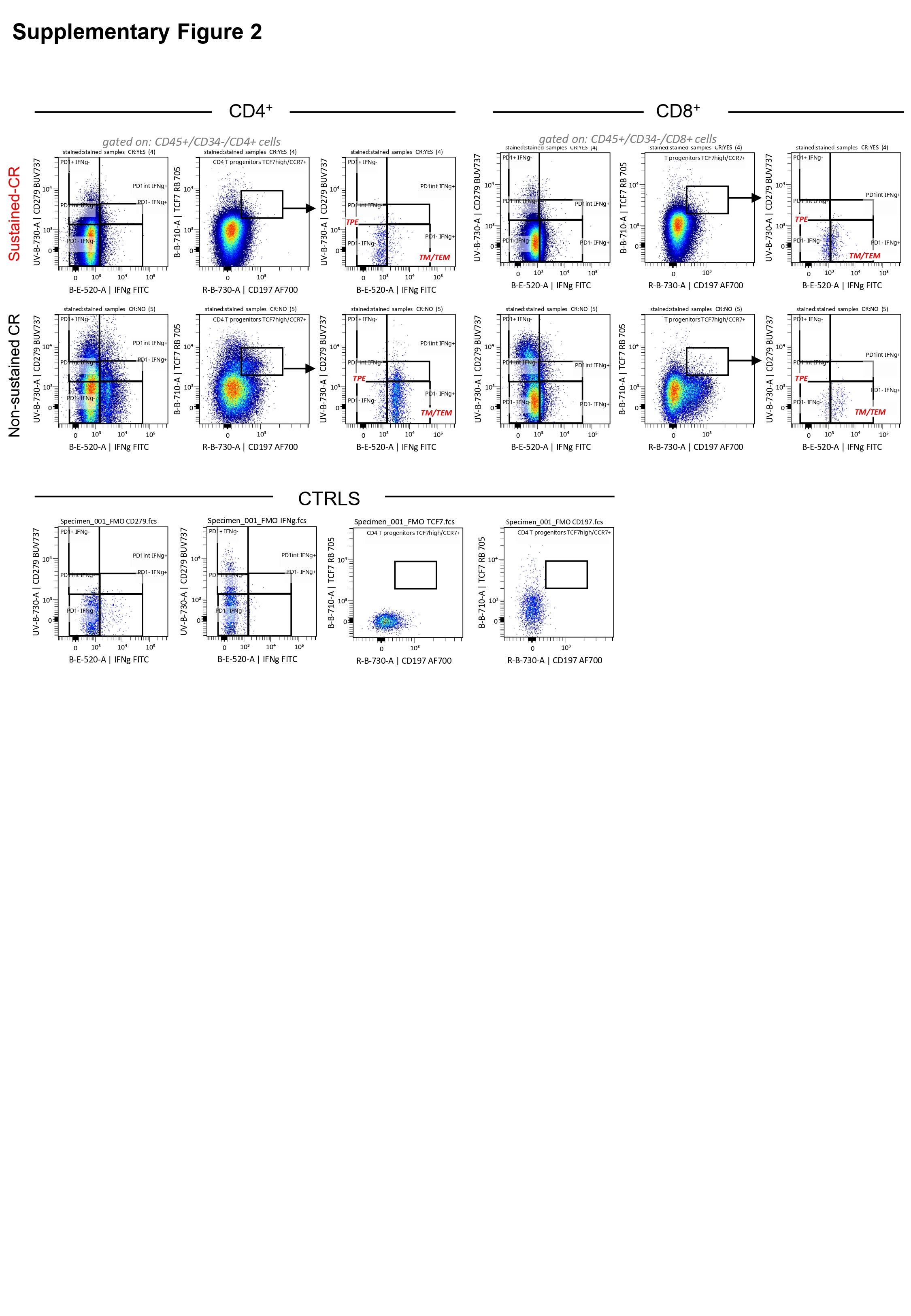

### Supplementary Figure 3.jpg

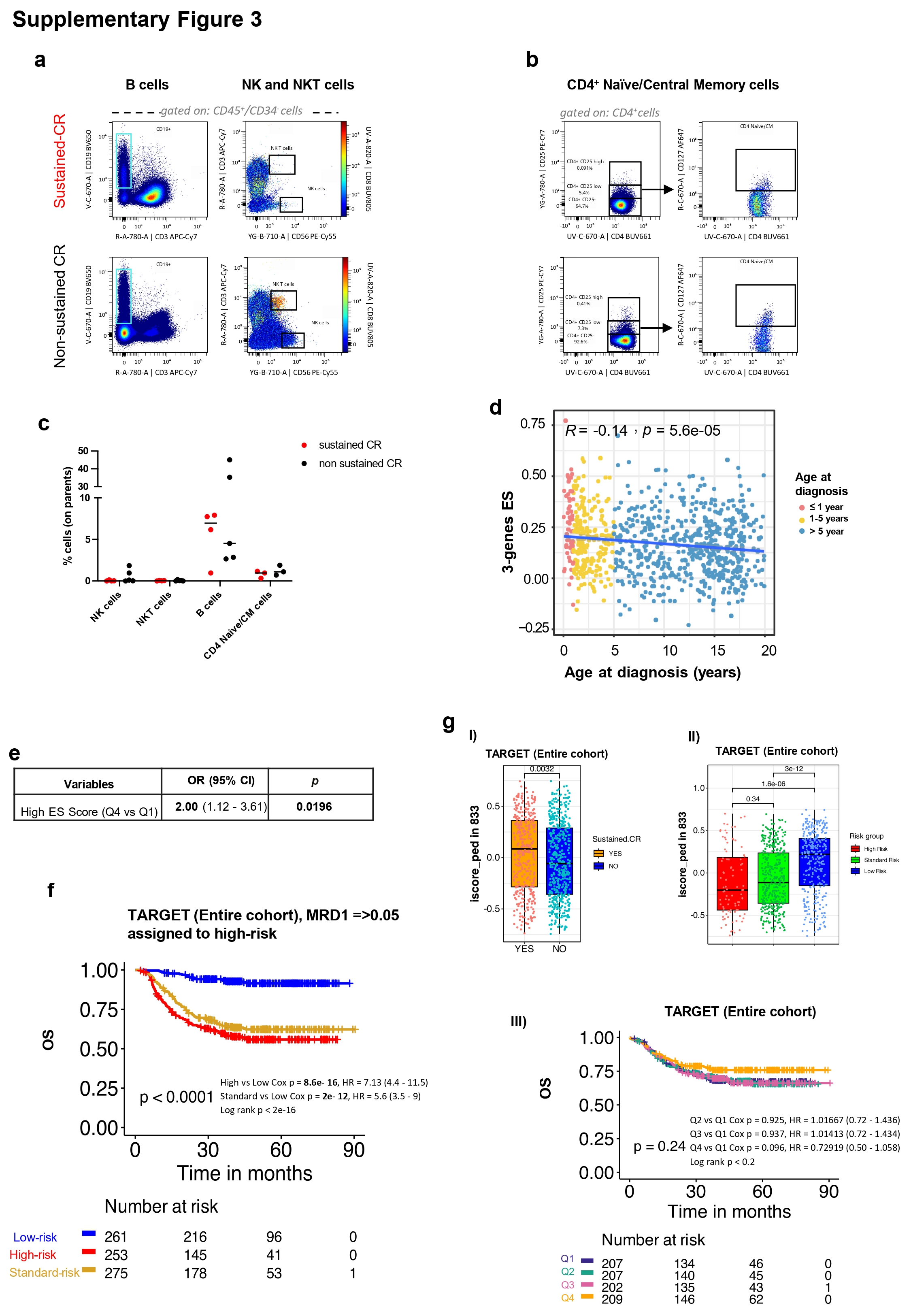
